# Tunable neuronal microenvironments drive distinct functional phenotypes in human iPSCs-derived dopaminergic neurons

**DOI:** 10.64898/2026.08.28.744416

**Authors:** Bahaa Daou, Lara Rodriguez, Paola Sartori, Gisela Luque, Maurizio Prato, Nuria Alegret, Sonia Alonso-Martín

## Abstract

Neuronal heterogeneity is a defining feature of complex neural circuits, where local differences in firing patterns, activity levels, and temporal dynamics shape information processing and emergent network activity. In adult neurons, this heterogeneity arises from a variety of known and unknown factors including the extracellular environment. Although neurons have been cultured on three-dimensional substrates, the effect of the microenvironments on their firing activity remains poorly understood. Here, we have synthesized two chitosan hydrogel systems seeded with human iPSCs-derived dopaminergic neurons as tunable platforms to control neuronal microenvironments. Both systems were formulated with the ability to incorporate carbon nanotubes (CNT), thus promoting neural interfacing. Calcium imaging combined with computational single-cell analysis demonstrated that supramolecular organization, hydration state, and general physicochemical properties differentially bias neuronal firing dynamics and synchrony leading to the emergence of distinct activity phenotypes despite identical cellular origin. These activity profiles were clustered through K-means and assigned to specific phenotypes including bursting irregular neurons, regular network contributors, or less active/quiescent neurons. Furthermore, CNTs incorporation enhanced local hydrogel compaction, resulting in unique active neuronal phenotypes, highlighting the potential of CNTs to modulate local cellular microenvironments. These findings establish tunable biomaterials as microenvironment contenders for controlling neuronal network state while giving insights on the interplay of different cues in promoting neuronal heterogeneity and functional phenotype relevant to neurodevelopment, neurodegeneration, and disease modelling.

## Introduction

In the central nervous system (CNS), neurons exhibit substantial functional heterogeneity, displaying diverse firing patterns and activity levels that collectively dictate information processing, plasticity, and tissue homeostasis.[1–3] Various extrinsic and intrinsic cues shape neuronal diversity, resulting in an estimated 10,000 distinct neuronal types with each neuronal subtype expressing a unique combination of ion channels, conferring specific excitation thresholds and firing patterns that contribute to the functional complexity of neural networks.[1] Indeed, complex circuitry give rise to diverse neuronal functions rather than relying on individual cell firing alone. [4]

The emergence of such heterogeneous activity states is a fundamental feature of neural development and although some factors governing their formation are known, many remain incompletely understood.[5–7] Although tissue engineering has predominantly focused on reproducing the intrinsic heterogeneity of tumors microenvironments, elucidating the role of the neuronal microenvironment in shaping neuronal activity and network maturation is essential for advancing our understanding of neurodegenerative diseases, guiding regenerative strategies, and developing physiologically relevant *in vitro* models.[8,9]

Due to the difficultness of establishing human primary neuronal cultures, the introduction by Takahashi *et al.* in 2007 of human induced pluripotent stem cells (hiPSCs) became a powerful platform for generating patient-specific neuronal models *in vitro* and understanding disease progression. [10–12] In particular, significant improvements in the differentiation of hiPSC-derived dopaminergic neurons, which are highly relevant for applications such as *in vitro* brain models and Parkinson’s disease (PD) research, have fueled growing interest in their use for both disease modelling and regenerative medicine.[13–15] Culturing iPSCs on top of tridimensional substrates increases their physiological relevance given the spatial geometry of native neuronal tissue, allowing the study of 3D network formation; however, the focus remains largely on phenotypic cues such as neuronal markers expression, adhesion, neuronal excitability and neurite outgrowth.[16–19] Despite the valuable insights these parameters give, functional characterization of neuronal populations and their similarity to the native-tissue remain limited. In contrast, both previous and current trends to understand CNS-related diseases, as well as, neuronal plasticity rely on functional assessment of neuronal networks of established complex networks such as brain-tissue-derived networks,[20,21] or organoid models rather than polymeric matrices.[22–25] Indeed, current computational tools employed to study these complex neuronal networks, whether in animal models or *in vitro* organoids, focus primarily on intrinsic individual or collective firing patterns derived from microscopic activity. [19–22] However, they fall short in identifying organizational clusters of neuronal phenotypes as a function of microenvironmental variations and the interplay between these factors in shaping distinct patterns of neuronal activity. [19–22] Moreover, growing evidence further suggest the existence of multiple functional states that coexist within developing neural networks.[20] Such heterogeneity may reflect differences in maturation, connectivity, excitability, or more importantly local microenvironmental influences including those apported by the extracellular matrix (ECM). The ECM plays a central role in regulating neuronal behavior by providing structural support and biochemical and biophysical signals that influence cell adhesion, neurite extension, synapse formation, and network development, which effectively can give rise to different microenvironment cues that would ultimately affect neuronal function and firing phenotype.[26–31]

Consequently, hydrogel-based biomaterials have emerged as attractive platforms for engineering neural microenvironments that better mimic native tissue conditions than conventional coated culture substrates.[32] Indeed, recent studies show that through the modulation of mechanical, structural, and chemical properties, hydrogels can actively influence cellular behavior and neuronal development.[33] Hence, employing polymeric scaffolds could facilitate studying the interplay of the microenvironment on neuronal activity and firing patterns in synthetic substrates.

Among the various polymeric materials investigated, chitosan, a polysaccharide obtained through the alkaline deacetylation of chitin, has gained considerable momentum since the 1980s, with particular focus on applications in neural regeneration and wound healing due to its biocompatibility and chemical properties.[34,35] Their major limitation in other application settings lies in their inherent degradability. [36] Although this property has been widely regarded as advantageous in tissue engineering due to the reported biocompatibility and bioactivity of its degradation products, it can compromise long-term structural integrity where prolonged scaffold stability is required.[34,37–39] Notably, these hydrogels often start to degrade within 7 days, limiting their use in *in vitro* studies.[40–42] To improve their stability, various strategies, including crosslinking, blending with other polymers, and tuning the pH of the medium, have further expanded their use *in vivo*. [43–46] Overall, the stability of chitosan-based hydrogels reported for *in vivo* applications, where prolonged structural stability is a critical design requirement, remains limited, with most scaffolds maintaining their integrity for 60-100 days before degrading, [43–46] Hence, their use as *in vitro* neuronal platforms or disease models remained scarce.

On the other hand, carbon nanotubes (CNT) have attracted considerable interest as hydrogel additives for neural interfacing. [47–50] Although the specific mechanisms remain unclear, CNTs have been shown to enhance synaptic activity, neural maturation, and neuronal specification through the modulation of cell signaling pathways, including the focal adhesion kinase (FAK)-AKT-β-catenin axis.[51–53] The major challenge remains the chemical modification and crosslinking of CNT-loaded chitosan hydrogels to enhance stability and avoid hydrogel degradation and swelling, while maintaining tuneability, elasticity and biocompatibility. Notably, using CNTs to induce higher neuron-material interaction while preserving the ability to tailor the hydrogel’s structure and mechanics provides an opportunity to investigate how defined microenvironmental conditions influence neuronal development and function. CNT-chitosan materials have previously been explored for neural cell culture and tissue engineering applications, yet their role in shaping functional neuronal activity patterns and network heterogeneity as a function of different mechanical properties remains largely unexplored. [36,54]

In the present study, we developed two tunable chitosan-based hydrogel systems with outstanding stability capable of incorporating CNTs and evaluated their potential as suitable microenvironments for the differentiation and culture of hiPSC-derived dopaminergic neurons. Using calcium imaging coupled with computational analysis of spontaneous activity, we investigated how variations in supramolecular organization, hydration state, and CNTs incorporation influence the emergence and organization of functional neuronal states. By linking hydrogel physicochemical properties with neuronal network behavior, this work provides new insight into the design of biomaterials capable of modulating functional neuronal phenotypes.

## Experimental Section

### Materials and reagents

Multi-walled CNTs (MWCNT, >95%) were purchased from Nanoamor Inc. (stock# 1237YJS: inner diameter, 5-10 nm; outside diameter, 20-30 nm; length, 0.5-2 μm), High molecular weight Chitosan CAS 9012-76-4 and 4-Amino benzoic acid (PABA) CAS 150-13-0 were purchased from Sigma-Aldrich. Phosphate-buffered saline (PBS) buffer was purchased in tablets and prepared following manufacturer procedures (Sigma-Aldrich), corresponding to 10 mM phosphate buffer containing 137 mM NaCl and 2.7 mM KCl at pH 7.3. The solvents were acquired from Carlo Erba Reagents SAS, Sabadell, Spain. All reagents and solvents were used as received with no further purification.

### Chitosan Hydrogel synthesis

Hydrogels were synthesized through a two-step process consisting of the maleoylation of chitosan followed by thermal crosslinking with PABA. Two hydrogel formulations, designated SL (for sponge-like; i.e.) and GL (for gel-like) (see below), were prepared by varying the thermal processing conditions while maintaining the same precursor composition. Corresponding CNT-containing hydrogels (SL-CNT and GL-CNT) were produced by incorporating MWCNTs into the precursor solution before gelation.

### Maleoylation of chitosan

Maleoylated chitosan (MCS) was synthesized by reacting chitosan (CS) with maleic anhydride. Briefly, 6 g of chitosan was dissolved in 100 mL of dimethyl sulfoxide (DMSO), followed by the addition of 3.5 g of maleic anhydride. The reaction mixture was maintained at 60 °C for 24 hours (h) under continuous stirring. Subsequently, the pH was adjusted using saturated sodium bicarbonate (NaHCO₃) solution until carbon dioxide evolution ceased, indicating complete neutralization of the excess maleic anhydride. The reaction mixture was dialyzed against Milli-Q water for 3 days with frequent water replacement to remove residual reagents and solvent. The purified product was then freeze-dried to obtain MCS as a dry powder.

### Hydrogel preparation

Hydrogels were prepared by dissolving 6% (w/v) of MCS, in deionized water at 80 °C for 30 minutes (min). 1.5% w/v of PABA was subsequently added, and the mixture was gently stirred at 80 °C for an additional 30 min while minimizing mechanical agitation to avoid premature gel formation. For CNT-containing formulations, MWCNTs (5 wt%, relative to the polymer weight) were first dispersed by sonication and subsequently incorporated into the MCS mixture before adding PABA under gentle stirring to obtain a homogeneous suspension. The resulting precursor solutions were processed using one of two thermal protocols:

- **SL hydrogels** were obtained by transferring the precursor solution directly into sealed reaction vials and incubating it at 37 °C for 7 days. Gelation was accompanied by syneresis, resulting in the expulsion of excess water from the hydrogel network. **GL hydrogels** were obtained by first heating the precursor solution at 100 °C for 3 h, followed by incubation at 37 °C for 7 days to complete gelation.

### Thermal Gravimetric Analysis (TGA)

Thermogravimetric analysis (TGA) was performed using a TGA Discovery (TA Instruments) under a nitrogen atmosphere with a flow rate of 25 mL·min⁻¹. Samples were equilibrated at 100 °C for 20 min and subsequently heated from 100 to 800 °C at a heating rate of 10 °C·min⁻¹. At least two independent measurements were performed for each sample. The resulting thermograms were analyzed using TRIOS software (version 4.4.0.41128, TA Instruments).

### Fourier Transform Infrared Spectroscopy (FTIR)

FTIR was performed on a ThermoScientific FTIR spectrometer in attenuated total reflectance (ATR) mode at room temperature (RT). FTIR spectra were recorded by accumulation of 64 scans in the range of 400 cm^-1^ till 4000 cm^-1^and a resolution of 4 cm^-1^.

### Swelling

Hydrogel discs (6 mm in diameter) were immersed in Milli-Q water immediately after synthesis for at least 72 h to reach swelling equilibrium. The swollen hydrogels were gently blotted to remove excess surface water and weighed to obtain the swollen weight (W_s_, g). Samples were then frozen at -20 °C for 24 h, followed by overnight storage at -80 °C, before being lyophilized using an Alpha 2-4 LSCplus freeze dryer (Part No. 102142) at a condenser temperature between -80 and -92 °C under a vacuum of 0.16 Pa. After complete drying, the samples were weighed again to determine the dry weight (W_d_, g). This stepwise freezing protocol was adopted to minimize frost shock and preserve the structural integrity of the hydrogels during freeze-drying.

The equilibrium swelling ratio was calculated according to:

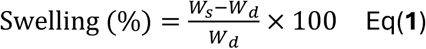

Subsequently, the freeze-dried hydrogels were immersed in PBS to simulate physiological conditions and incubated at 37 °C. Samples were removed after 2, 4, 24h, as well as at later time points up to 2 years, gently blotted to remove excess surface liquid, and weighed. The swelling ratio at each time point was calculated using the equation above.

### Rheology analysis

Frequency sweep measurements were performed on all hydrogel samples pre-cut into discs (8 mm in diameter) using an MCR 301 rheometer (Anton Paar) equipped with an 8 mm patterned parallel-plate geometry. Measurements were carried out at RT at a constant strain of 0.016%. Prior to the frequency sweep, amplitude sweep tests were performed to determine the linear viscoelastic region (LVR), defined as the range in which both the storage modulus (G′) and loss modulus (G″) remain independent of the applied strain at a fixed frequency of 10 Hz.

### Nuclear Magnetic Resonance (NMR) imaging

The degree of maleoyl substitution of MCS was determined by proton nuclear magnetic resonance (^1^H NMR) spectroscopy using a 500 MHz Bruker spectrometer (Bruker BioSpin, Germany). MCS was dissolved in D₂O at a concentration of 10 mg·mL⁻¹ with heating to ensure complete solubilization. Pyridine (2.6 mg·mL⁻¹) was added as an internal standard for quantitative analysis. The degree of substitution was determined by integrating the characteristic maleoyl proton signals relative to the pyridine resonance, as described in the supplementary information (**Figures S1-S2 and Table S1**). All spectra were recorded at 25 °C.

### Scanning Electron Microscopy (SEM)

The microstructure of the hydrogels was characterized using a field-emission SEM (FE-SEM; JSM-IT800HL, JEOL, Tokyo, Japan) equipped with a field emission gun (FEG), Beam Deceleration (BD) stage-bias technology, and secondary electron (SE), backscattered electron (BSE), and transmitted electron detectors. Hydrogel samples were cut into cylindrical specimens (3 mm in diameter), shock-frozen in liquid nitrogen, and subsequently lyophilized prior to imaging. SEM images were acquired in high-vacuum mode without conductive sputter coating using accelerating voltages ranging from 2.5 to 5.0 kV.

### Mechanical properties and Young’s Modulus (YM)

Uniaxial compression tests were performed using a universal testing machine (Instron) at a crosshead speed of 10 mm·min⁻¹ to obtain stress-strain curves. Hydrogel discs (6 mm in diameter) were prepared using a biopsy punch, and their thickness was measured prior to testing using digital micrometer. Following the initial compression test, the samples were rehydrated and subjected to a second compression test under identical conditions. The YM was determined for both the initial compression and recompression from the slope of the initial linear region of the corresponding stress-strain curves. Tests were performed in triplicates.

### Differentiation of hiPSCs into ventral midbrain dopaminergic neurons

hiPSCs were differentiated into ventral midbrain dopaminergic (vmDA) neurons following a modified version of the chemically defined, xeno-free protocol reported by Gantner *et al.[13]* Briefly, hiPSCs during early passages phases (P1-P5) were cultured on top of pre-cut hydrogels (Seeding density: 500K, ca. 6 mm x 3 mm hydrogels) under feeder-free conditions after which differentiation was initiated using a dual-SMAD inhibition strategy. Briefly, *[13]* neural induction was achieved in N2B27 induction medium supplemented with 10 μM SB431542 and 200 nM

LDN193189 to inhibit the TGF-β and BMP signaling pathways, respectively. Ventral midbrain patterning was induced through the addition of 100 ng·mL⁻¹ Sonic Hedgehog (SHH) and 2 μM purmorphamine from days 1 to 5, while activation of Wnt signaling was achieved using 2.5 μM CHIR99021 from days 2 to 11 to promote caudalization toward a midbrain dopaminergic fate. SB431542 was withdrawn after day 5, whereas LDN193189 was maintained until day 11. On day 11, cultures were transitioned to maturation medium containing 20 ng mL⁻¹ brain-derived neurotrophic factor (BDNF), 20 ng·mL⁻¹ glial cell line-derived neurotrophic factor (GDNF), 0.1 mM dibutyryl cyclic AMP (dbcAMP), 200 μM ascorbic acid, 1 ng·mL⁻¹ transforming growth factor-β3 (TGF-β3), and 10 μM DAPT to promote neuronal maturation and dopaminergic specification. Cells were maintained under standard culture conditions (37 °C, 5% CO₂) with medium changes every other day until day 30, when they were fixed with 4% paraformaldehyde (PFA) followed by cold PBS washes 3 times. Two different set of experiments were carried out for immunofluorescence and calcium imaging, within each, triplicates of each hydrogel were used.

### Immunofluorescence

After fixation, samples were stored at 4 °C until staining. A blocking/permeabilization solution consisting of 0.25% Triton X-100 in Dulbecco’s (D)PBS supplemented with 2.5% fetal bovine serum (FBS) and 2.5% normal goat serum (NGS) was prepared. Samples were incubated in the blocking solution for 2 h using a volume sufficient to fully immerse the hydrogels, which was adjusted according to their swelling behavior. An antibody dilution buffer was then prepared by diluting the blocking solution 1:10 in DPBS. Primary antibodies to identify the neuronal markers, guinea pig anti-NeuN (nuclear/perinuclear protein encoded by *RBFOX3*) and rabbit anti-TUBB3 (βIII-tubulin, major component of neuronal microtubules), were diluted 1:1000 in the antibody dilution buffer and added at a volume equivalent to six times the hydrogel volume. Samples were incubated overnight at 4 °C. Following primary antibody incubation, samples were washed three times with DPBS using gentle pipetting to avoid disrupting the hydrogels. Secondary antibodies, goat anti-rabbit Alexa Fluor 488 and goat anti-guinea pig Alexa Fluor 647, were diluted 1:500 in antibody dilution buffer and incubated with the samples for 2 h using the same antibody-to-hydrogel volume ratio. Samples were subsequently washed three times with DPBS before incubation with 4′,6-diamidino-2-phenylindole (DAPI) for 1 h for nuclei staining, followed by three additional DPBS washes. Finally, the samples were transferred to glass-bottom imaging dishes (#1.5H coverslip thickness) and imaged using a laser scanning confocal microscope (LSM 900, Zeiss).

### Images acquisition and quantification

Maximum-intensity Z-projections were generated from confocal image stacks using ZEN 2.3 software (Zeiss). Fluorescence image analysis was performed in ImageJ (National Institutes of Health, USA; Java version 13.0.6). NeuN and TUBB3 fluorescence signals were automatically segmented by intensity thresholding from their respective fluorescence channels. Because the hydrogels exhibited autofluorescence within the 410 nm spectral region, a binary mask was generated from the overlap of the DAPI (UV light excited) and TUBB3 (Alexa Fluor 488) channels to identify the hydrogel area, which was subsequently excluded from the quantitative analysis. The fluorescence-positive areas corresponding to NeuN and TUBB3 were measured, and the ratio F_NeuN_/F_TUBB3_ was calculated as an indicator of neuronal maturation. In addition, the fluorescence-positive area of each marker was normalized to the nuclear stained DAPI-positive area and expressed as F_NeuN_/_DAPI_ and F_TUBB3_/_DAPI_.

Quantitative analysis was performed on three independent regions of interest (ROIs), each measuring ca. 1 mm x 1 mm, per replicate. Data are presented as the mean ± standard deviation (SD) of triplicate measurements. Outliers were identified using the ROUT method (Q = 5%) and confirmed with Grubbs’ test (α = 0.05). Prism GraphPad (Version 11.0.2) was used for statistical analysis. One outlier was detected in the SL hydrogel group for F_NeuN_/F_TUBB3_ analysis and two outliers were detected in the SL group for F_NeuN_/_DAPI_ and F_TUBB3_/_DAPI_ analyses. values were excluded from subsequent analyses. Statistical comparisons among groups were performed using Kruskal-Wallis test followed by Dunn’s multiple-comparisons test, with statistical significance defined as *p* < 0.05.

### Calcium imaging

Following differentiation, the culture medium was aspirated from wells containing hydrogel-seeded hiPSCs, and the samples were gently washed once with PBS to remove residual culture medium that could interfere with fluorescence measurements. Intracellular calcium was monitored using the fluorescent calcium indicator FLUO-8 (ab112129, Abcam) according to the manufacturer’s instructions. Briefly, cells were incubated with the FLUO-8 loading solution for up to 1 h at 37 °C to allow intracellular dye loading, de-esterification, and equilibration prior to image acquisition. To minimize sample manipulation and preserve the integrity of the hydrogel-cell constructs, calcium imaging was performed directly in the original culture wells using an Axio Observer inverted epifluorescence microscope (Zeiss, Germany) equipped with a motorized stage, environmental incubation chamber, and Zen Blue software (v2.3). Fluorescence images were acquired using the 38 HE eGFP filter set (excitation 450-490 nm, dichroic beamsplitter 495 nm, emission 500-550 nm). Images were recorded using the monochrome Axiocam MR R3 camera and along 2000 frames corresponding to around 1000 s time frame. Image sequences were analyzed using ImageJ. Background fluorescence was measured for each ROI and subtracted from the corresponding fluorescence traces to generate background-corrected time series. These corrected fluorescence traces were subsequently used for computational calcium transient analysis and quantitative data interpretation.

### Computational calcium imaging workflow

Calcium imaging datasets were processed using a custom computational workflow to quantify neuronal activity and identify functional activity patterns across experimental groups. Fluorescence intensity traces were extracted for each manually defined ROI corresponding to individual neurons. Raw fluorescence signals were cleaned by removing empty values and converted to numerical arrays prior to analysis. For each neuron, the fluorescence baseline (F₀) was estimated as the 10th percentile of the fluorescence trace according to the following equation:

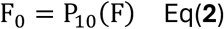

where *P*_10_ denotes the 10th percentile of the fluorescence trace. Normalized fluorescence changes were calculated as:

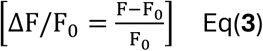

where *F* is the fluorescence intensity at each time point and *F₀* is the estimated baseline fluorescence. Normalized traces were subsequently smoothed using a Gaussian filter (σ = 1) to reduce high-frequency noise while preserving transient dynamics. Calcium transients were detected automatically using an adaptive threshold based on the standard deviation of each normalized trace. Peaks exceeding 2.5 standard deviations above baseline and persisting for a minimum duration of two consecutive frames were classified as calcium events. For each detected event, a temporal window surrounding the transient was extracted and used for subsequent feature analysis. Feature standardization was performed before clustering neurons using K-means algorithm which minimizes the Euclidean distance between x and c according to the following equations:

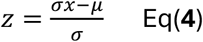

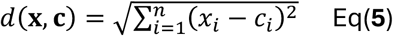

x is the feature vector of a neuron, c is the centroid of a cluster, *x_i_* is the *i*-th feature of the neuron, *c_i_* is the corresponding centroid value, *n* is the total number of extracted features. Silhouette analysis was carried out for the determination of optimal cluster number:

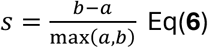

However, manual clustering K number was pre-assigned followed by analyzing the data of raster plot, etc., to verify significance and effectiveness of the cluster number. Cluster assignments were subsequently used to generate raster plots, activity heatmaps, and average activity traces representing the temporal dynamics of each functional neuronal population. Neuronal activity was evaluated by calculating the coefficient of variation (CV) and time-resolved CV to determine true variability within a population through the duration of acquisition described by the following equation:

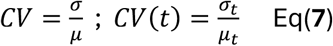

whereby, *σ* is the standard deviation and *μ* is the mean Δ*F*/*F*_0_ signal for each neuron.

The previous info was then analyzed and for experimental groups presenting a continuum and heterogenous firing pattern, mean neuronal activity was additionally calculated to those calcium transients, thereby providing a more representative estimate of functional network activity.

## Results and Discussion

### Physicochemical properties regulate the mechanical and structural microenvironment

To establish the structure-function relationship governing dopaminergic neuronal maturation, chitosan-based hydrogels were engineered through a two-step supramolecular strategy and systematically characterized prior to biological evaluation. Maleoylation of chitosan introduced pendant carboxyl groups capable of increasing intermolecular interaction sites within the polymer backbone (**Figure 1A**). Subsequent crosslinking with PABA generated two hydrogel formulations with distinct supramolecular network organizations: SL, which underwent syneresis during synthesis and formed a lower-water-content network, and GL, which retained a substantially more hydrated architecture. CNTs were subsequently incorporated into both formulations to generate SL-CNT and GL-CNT. Successful maleoylation was confirmed by the appearance of a characteristic FTIR band at 1708 cm⁻¹, corresponding to the C=O stretching vibration of the introduced carboxyl groups (**Figure S3A**). This maleolation modified the pH responsiveness of chitosan, rendering MCS soluble under neutral and alkaline conditions, consistent with previous reports.[55] FTIR further suggested that supramolecular crosslinking was predominantly mediated through hydrogen bonding, as evidenced by shifts in the N–H bending vibration (1550 cm⁻¹) and the symmetric COO⁻ stretching vibration (1371 cm⁻¹). Successful functionalization was further confirmed by NMR and thermogravimetric analysis (**Figures S1,S2,S3B and Table S1**).

**Figure 1.**
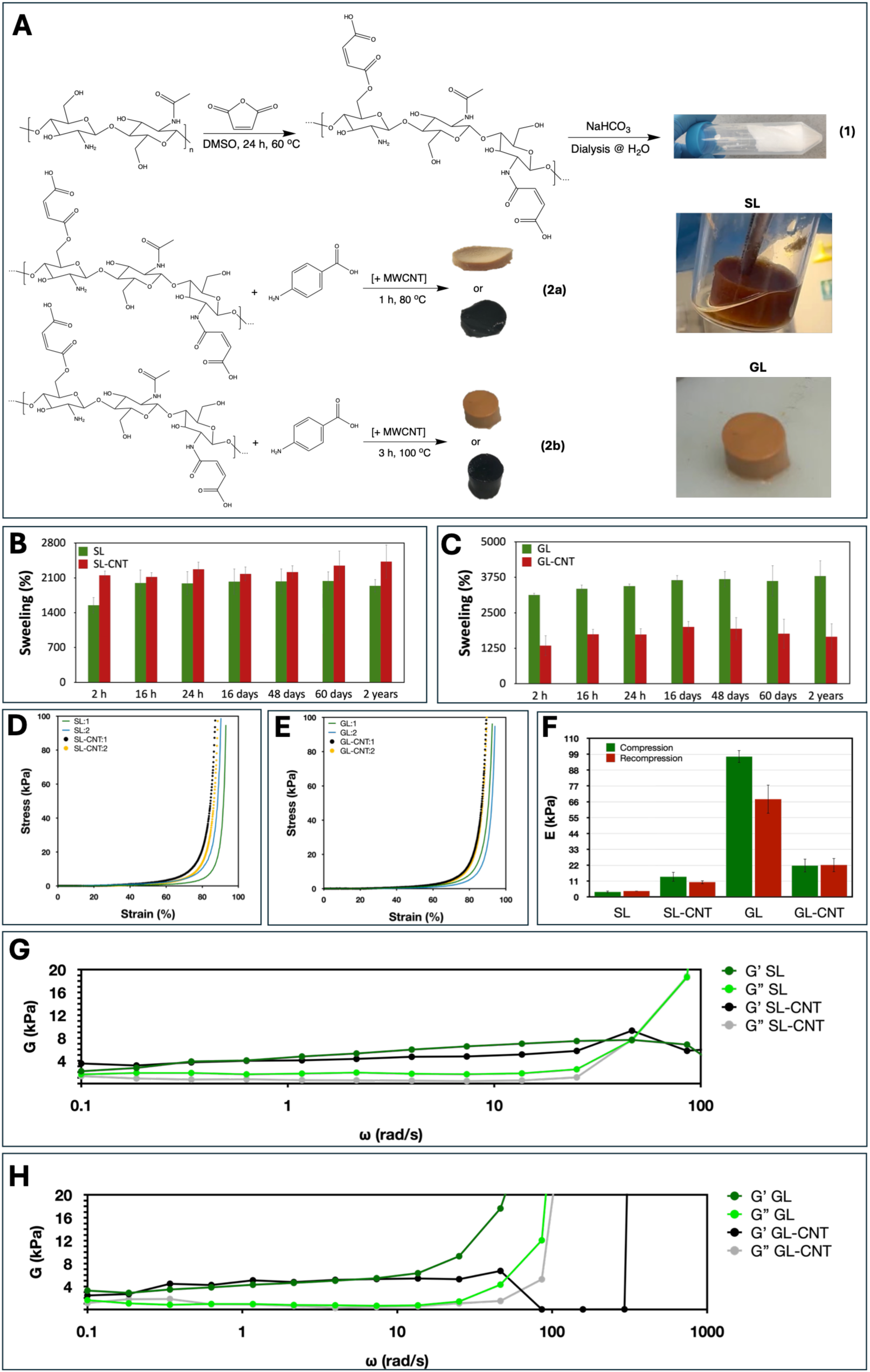
Physicochemical and mechanical characterization of SL and GL chitosan hydrogel systems. (**A**) Schematic illustration of the synthesis route used to prepare the SL and GL chitosan hydrogels. (**B,C**) Equilibrium swelling ratio of the SL and GL hydrogel systems, respectively, measured after immersion in PBS at 37 °C for 2 h up to 2 years. (**D,E**) Representative compressive stress–strain curves obtained for the SL and GL hydrogel systems, respectively at a crosshead speed of 10 mm·min⁻¹. (**F**) Young’s modulus (YM) of all hydrogel formulations, calculated from the initial linear region of the stress-strain curves. (**G,H**) Frequency sweep analysis showing the storage modulus (Gʹ) and loss modulus (Gʺ) of the SL and GL hydrogel systems, respectively, measured at 0.016% strain within the linear viscoelastic region determined by amplitude sweeps over a frequency range of 10 Hz at 21 °C. h, hours. Data are presented as mean ± SD. N = 3 independent samples.

The distinct supramolecular organizations were reflected in both the thermal stability and hydration behavior of the hydrogels. Thermogravimetric analysis revealed that GL consistently exhibited a higher residual mass than SL, both in the absence and presence of CNTs, suggesting a greater degree of intermolecular network organization (**Figure S3B**). [56] These differences were further reflected in the swelling behavior (**Figure 1B,C**). Both formulations rapidly reached equilibrium within approximately 2 h and maintained structural integrity under physiological conditions for two years at 37 °C, demonstrating excellent long-term stability. However, SL stabilized at an equilibrium swelling ratio of approximately 2,000%, whereas GL reached approximately 3,500%, reflecting a substantially greater capacity for water uptake upon rehydration. This difference likely reflects the distinct balance between polymer–water interactions and network constraints in the two supramolecular architectures.

Interestingly, CNTs incorporation affected the two systems differently. While SL-CNT exhibited minimal changes in swelling relative to SL, GL-CNT showed a marked reduction to approximately 1,700%, indicating stronger CNT-induced hydrophobic interactions. Importantly, these measurements reflect the ability of the hydrogels to re-swell after freeze-drying rather than their absolute hydration state. Thus, the swelling ratio should not be interpreted as directly proportional to the water content of the original gels. Because the SL system undergoes syneresis, it retains a lower hydration state than GL during gel formation, both with and without CNT incorporation. Thus, the reduced re-swelling capacity of GL-CNT does not necessarily imply a lower native water content than that of SL-CNT (**Figure 1B,C**). Consistent with these observations, SEM images showed that the GL hydrogels exhibited a more compact porous microstructure compared to SL hydrogels (**Figure S4**).

To further probe differences in molecular organization, solid-state fluorescence spectroscopy was performed (**Figure S3C,D**). Both SL and GL exhibited broadened and red-shifted emission spectra compared with pristine chitosan and MCS, while GL consistently showed greater spectral broadening than SL, supporting a higher degree of intermolecular organization. In contrast, MCS displayed a pronounced blue shift in excitation accompanied by a red shift in emission relative to pristine chitosan, producing a substantially larger Stokes shift (**Figure S3C,D**). This distinct photophysical response suggests that maleoylation alters the excited-state relaxation pathway, consistent with the formation of additional or modified emissive states within the functionalized polymer backbone. Although hydrogen bonding appears to be the dominant crosslinking mechanism, additional contributions from π–H interactions and localized PABA crystallization cannot be excluded and may further reinforce the supramolecular network. [57,58]

Collectively, the FTIR, TGA, fluorescence, and swelling analyses demonstrate that subtle modifications in supramolecular network organization produce hydrogels with markedly different hydration states while maintaining excellent physiological stability. These differences establish distinct physicochemical microenvironments that subsequently dictate the mechanical properties of the hydrogels and, ultimately, their ability to regulate neuronal maturation and function.

### Hydrogel network organization enables independent tuning of viscoelasticity and compressive mechanics

Given the distinct hydration states and supramolecular network organizations established above, we next investigated how these differences translated into the mechanical microenvironment experienced by differentiating neurons. All hydrogel systems exhibited a nonlinear, J-shaped compressive stress–strain response (**Figure 1D,E**), characterized by increasing resistance to deformation at higher strains. Such strain-stiffening behavior is seen in various biological tissues and is consistent with progressive network densification and increasing load-bearing interactions under compression, together with the contribution of fluid redistribution within the hydrated network.[59] All formulations sustained compressive strains approaching 80%, demonstrating large-deformation tolerance. Despite their common deformation profile, the two hydrogel systems displayed markedly different mechanical behaviors. The highly hydrated GL network exhibited a substantially higher YM (97.4 kPa) than SL (3.5 kPa) (**Figure 1F**). Notably, the greater stiffness of GL suggests that its enhanced supramolecular organization dominates its mechanical response, providing a more efficient load-bearing, while fluid redistribution within its highly hydrated porous matrix may additionally contribute to its compressive response. In contrast, the less organized SL network deformed more readily under compression, resulting in a significantly lower modulus.

Consistent with the swelling results, CNT incorporation affected the two hydrogel networks differently. SL-CNT exhibited the expected reinforcing effect of CNTs, increasing the YM from 3.5 to 13.9 kPa, while producing minimal changes in swelling. Conversely, GL-CNT displayed a pronounced reduction in stiffness from 97.4 to 21.7 kPa, suggesting that CNT incorporation reorganized or partially perturbed the highly associated supramolecular interactions responsible for the mechanical strength of GL rather than simply providing additional reinforcement. The simultaneous reduction in GL-CNT swelling and YM further indicates that CNT incorporation altered both water–network interactions and the organization of the load-bearing polymer network (**Figure 1F**). Thus, whereas CNT-mediated reinforcement appears to dominate in SL, CNT–matrix interactions have a substantially greater effect on the pre-existing supramolecular organization of GL.

These differences were further reflected in the recovery behavior following compression. SL and SL-CNT recovered their original mechanical properties after rehydration and recompression, displaying characteristic sponge-like recovery (**Figure 1F**). In contrast, GL exhibited an approximately 30% reduction in YM following recompression, indicating partial rearrangement of its supramolecular network during the initial loading cycle. Interestingly, CNT incorporation largely preserved the compressive response of GL-CNT following rehydration. Thus, although CNTs reduced the initial stiffness of GL, they also appeared to stabilize the reorganized network against further mechanical changes during subsequent compression cycles (**Figure 1F**).

Together, these results demonstrate that the hydrogel network organization governs not only the water content but also mechanical stress redistribution and elasticity. Importantly, the opposite responses of the SL and GL systems to CNTs incorporation indicate that reinforcement is governed by network architecture rather than by CNTs content alone, providing a versatile strategy for independently tuning the mechanical microenvironment presented to differentiating neurons.

### Microenvironment tuning give raise to different maturation states and neuronal clusters of hiPSC-derived dopaminergic neurons

hiPSCs were differentiated directly on each hydrogel formulation to investigate how the engineered microenvironment regulates neuronal maturation and function. Hydrogel composition markedly influenced neuronal morphology, NeuN and TUBB3 expression, neuronal network organization, and calcium firing dynamics. Both SL and GL supported the differentiation of mixed neuronal populations containing TUBB3+ and NeuN+ neurons together with NeuN- or TUBB3-cells, while displaying evident neurite extension throughout the hydrogel network (**Figure 2A**). However, the two microenvironments promoted distinct maturation profiles. Compared with SL, GL exhibited an almost two-fold increase in NeuN expression (**Figure 2B,C**), demonstrating that supramolecular organization and network hydration substantially enhance neuronal maturation while preserving neurite outgrowth. CNTs further modified this response. Consistent with previous reports,[60] CNTs incorporation promoted the development of the TUBB3+ neuronal network, with the strongest effect observed in the SL-CNT hydrogel, which displayed a statistically significant increase compared with SL. GL-CNT showed a modest, non-significant increase relative to GL. Because thin neuritic processes contribute relatively little to the total fluorescence-positive area, subtle differences in neurite extension may be visually apparent without producing a statistically significant change in area-based quantification. (**Figure 2B,D**). Importantly, despite the increase in TUBB3 expression following CNTs incorporation, GL-CNT maintained higher NeuN expression than its SL counterpart, preserving the maturation trend established by the hydrated and more supramolecularly organized GL microenvironment (**Figure 2B**). Interestingly, GL and GL-CNT also displayed a distinct spheroidal cellular organization, suggesting that enhanced cell-to-cell interactions within this environment may facilitate neuronal maturation and network assembly during differentiation (**Figure 2A; GL-CNT_zoomed_**).

**Figure 2.**
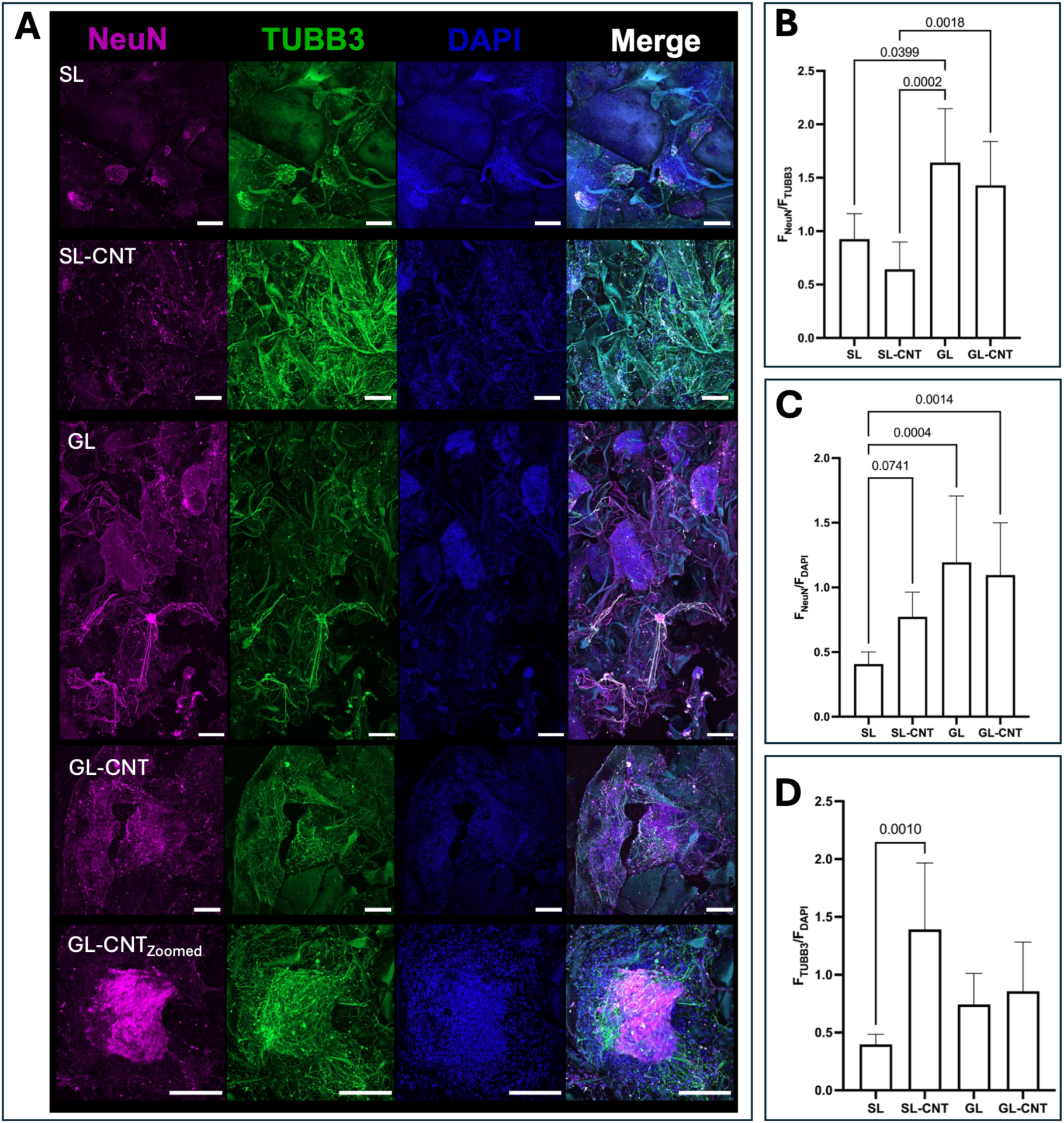
Immunofluorescence analysis of dopaminergic neurons following differentiation across hydrogel formulations. (**A**) Representative confocal fluorescence images of cells cultured within each hydrogel formulation and stained for NeuN (mature neurons, magenta), βIII-tubulin (TUBB3; neurons and neurites, green), and DAPI (nuclei, blue), together with the corresponding merged images. Scale bars represent 200 μm. (**B**) Quantification of neuronal maturation expressed as the ratio of NeuN to TUBB3 fluorescence intensity (F_NeuN_/F_TUBB3_). (**C**) NeuN fluorescence intensity normalized to DAPI fluorescence intensity (F_NeuN_/F_DAPI_). (**D**) TUBB3 fluorescence intensity normalized to DAPI fluorescence intensity (F_TUBB3_/F_DAPI_). Data are presented as mean ± SD from n = 3 independent hydrogel replicates per formulation. For each replicate, three regions of interest larger than 1 × 1 mm were imaged and analyzed. Statistical significance was assessed using the Kruskal–Wallis test followed by Dunn’s multiple-comparisons test. p < 0.05 was considered statistically significant.

To determine whether these structural differences were accompanied by differences in functional neuronal behavior, spontaneous calcium activity was analyzed using a computational pipeline that identified calcium events, classified neuronal activity patterns, and quantified population-level activity dynamics (**Figure 3**). Mean activity profiles first revealed clear differences in the extent of functional recruitment among the hydrogel formulations. In SL hydrogels, spontaneous calcium activity was restricted to only a limited fraction of the neuronal population, whereas the other hydrogel formulations displayed activity distributed across a substantially larger proportion of neurons (**Figure 3A-D**). These differences were also evident in the corresponding heatmaps (**Figure 3E-H**). The SL heatmap exhibited diffuse signal variation rather than discrete, high-intensity calcium transients (**Figure 3E**). In contrast, GL displayed several neurons with large ΔF/F₀ transients (**Figure 3F**), while both SL-CNT and GL-CNT exhibited prominent calcium signals distributed across larger fractions of the neuronal population (**Figure 3G,H**). Thus, CNT incorporation was associated with a further increase in neuronal activity and functional recruitment.

**Figure 3.**
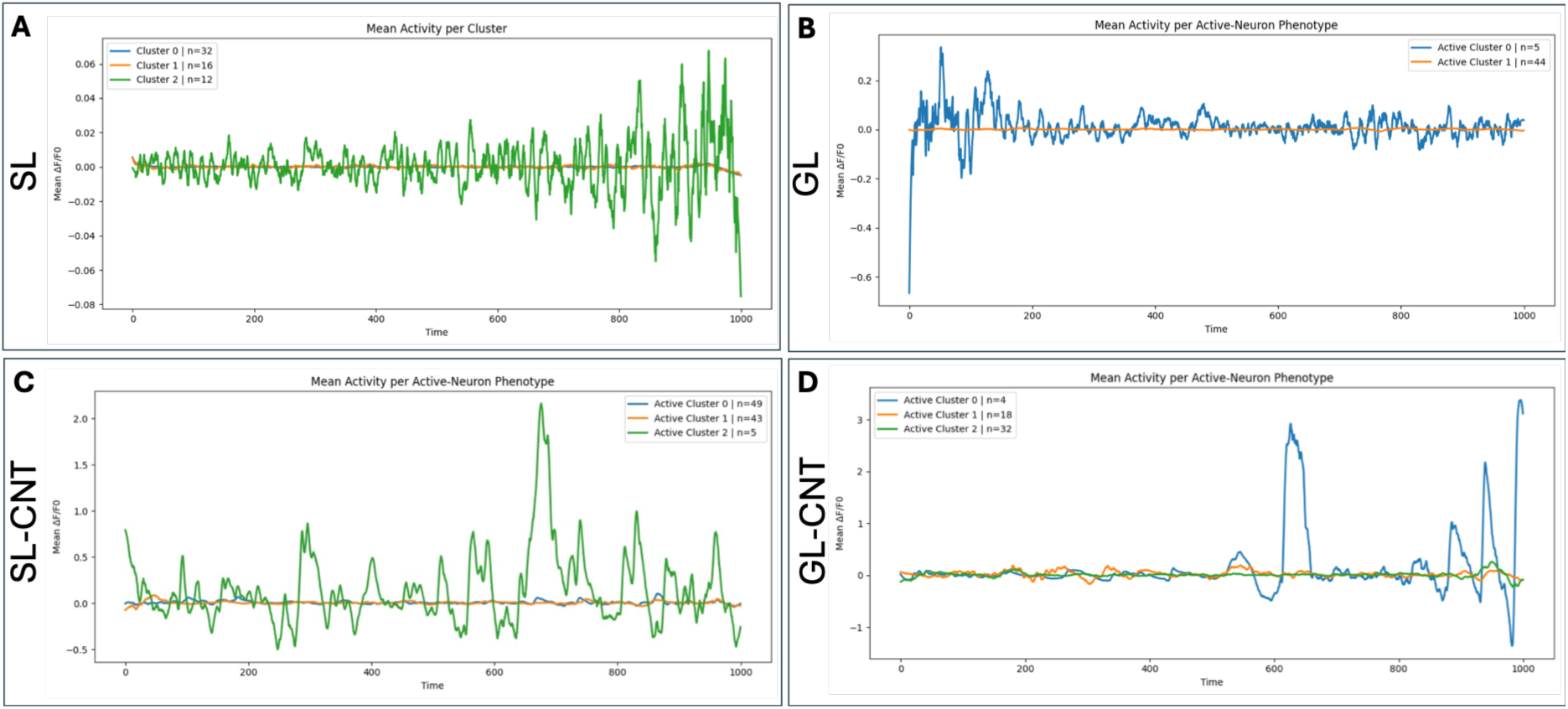
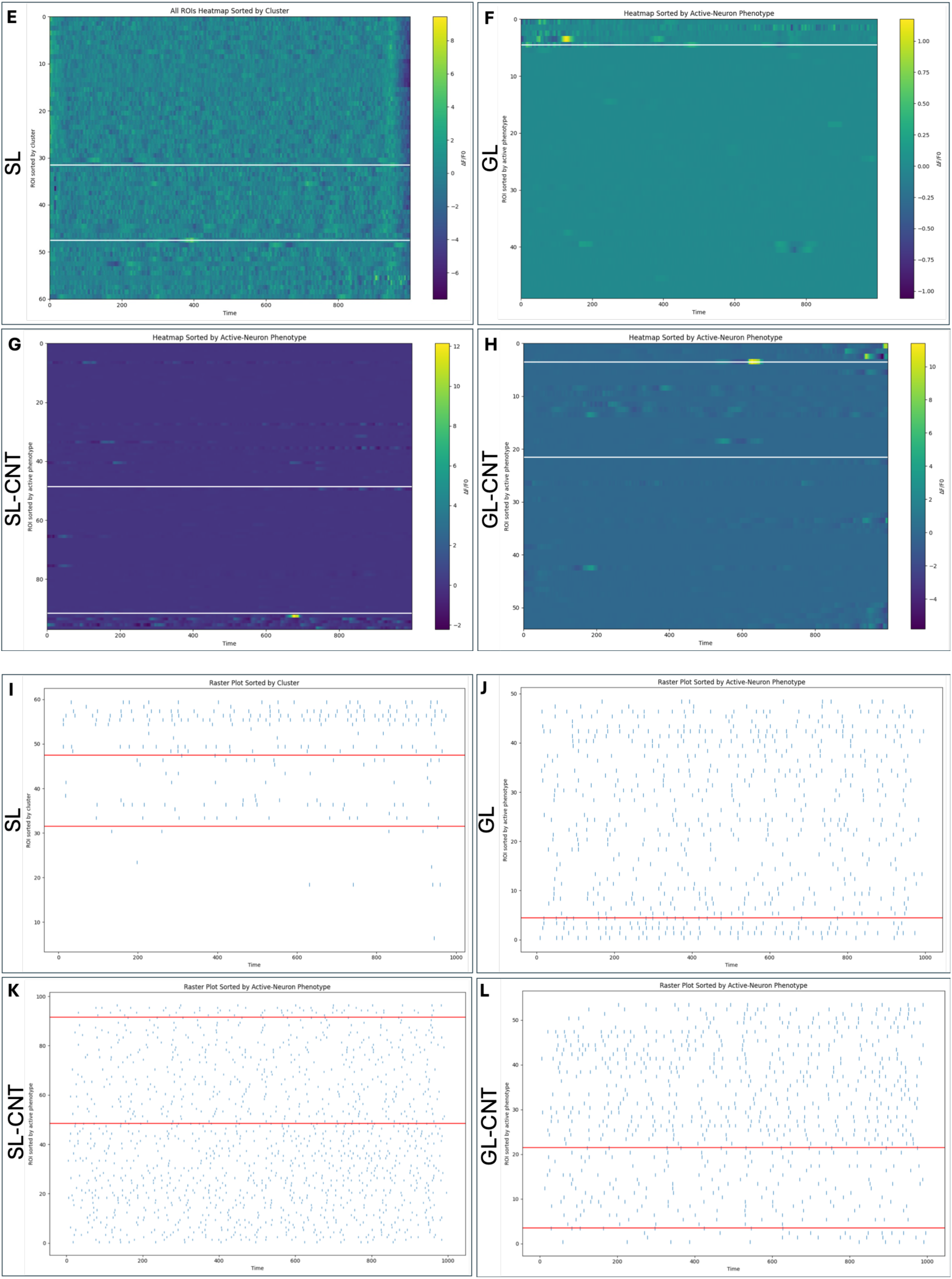
Computational analysis of distinct neuronal activity phenotypes across hydrogel formulations. (**A**) Mean calcium activity traces of the computationally identified neuronal activity clusters in the SL hydrogel. (**B–D**) Mean calcium activity traces of the computationally identified active neuronal phenotypes in the GL, SL-CNT, and GL-CNT hydrogels, respectively. (**E–H**) Heatmaps of normalized calcium fluorescence signals from individual neurons cultured in the SL, GL, SL-CNT, and GL-CNT hydrogels, respectively. Neurons in the SL hydrogel were grouped according to their assigned activity cluster, whereas active neurons in the GL, SL-CNT, and GL-CNT hydrogels were grouped according to their assigned active neuronal phenotype. (**I–L**) Corresponding raster plots showing the temporal firing patterns of neurons cultured in the SL, GL, SL-CNT, and GL-CNT hydrogels, respectively. Heatmaps display normalized ΔF/F₀ signals over time, with each row representing an individual neuron. Raster plots show detected calcium transients in binary format, with each event indicating the occurrence of a calcium transient. Neurons were grouped using the computational clustering workflow described in the Methods.

While mean activity profiles and heatmaps described the extent and magnitude of calcium activity, raster plots and activity-based clustering were used to resolve how this activity was distributed across distinct neuronal phenotypes (**Figure 3I-L**). In SL, the remaining neurons were predominantly silent (**Figure 3I**), and this less supramolecularly organized microenvironment was associated with three distinct activity clusters: a large population of silent neurons and two active phenotypes corresponding to regular contributors and highly active or bursting neurons. These observations were consistent with the reduced NeuN and TUBB3 expression observed in SL, together indicating that this microenvironment biases the neuronal population toward a less mature, predominantly silent functional state. In particular, microenvironments that maintain reduced neuronal maturation, limited functional recruitment, or network hypoactivity could provide useful platforms for modelling neurodevelopmental phenotypes characterized by impaired maturation or reduced neuronal activity.

Unlike SL, the other hydrogel formulations contained substantially larger active neuronal populations, making clustering based solely on overall activity magnitude less informative. Consequently, active neurons were classified according to their firing dynamics, revealing different functional phenotypes comprising sparsely active neurons, regular network contributors, and highly active or bursting neurons. GL was predominantly composed of regular contributors, together with a smaller population of highly active neurons (**Figure 3J**). The broader functional recruitment and predominance of a regular-contributor phenotype, together with the increased neuronal marker expression, were consistent with a more functionally mature and comparatively uniform active population.

Notably, despite differences in neuronal marker expression between the SL- and GL-based systems, CNTs incorporation appeared to promote a broadly similar distribution of active neuronal phenotypes. In both SL-CNT and GL-CNT, highly active or bursting neurons represented the largest active population, followed by a slightly smaller proportion of regular contributors and only a minor population of sparsely active neurons (**Figure 3K,L**). However, the largest calcium transients were not confined exclusively to a specific activity phenotype. In SL-CNT, neurons within the lower-event-count cluster displayed several high-amplitude calcium transients, while strong signals were also observed among highly active neurons (**Figure 3G,K**). This partial uncoupling between event frequency and calcium-transient amplitude is particularly interesting, as similar functional alterations have been reported in neurometabolic disorders such as succinic semialdehyde dehydrogenase (SSADH) deficiency,[61,62] in which reduced event frequency can coexist with markedly increased calcium-transient amplitudes. Thus, the SL-CNT microenvironment may provide a useful 3D setting for investigating neuronal phenotypes characterized by dysregulated coupling between activity frequency and calcium signaling. In contrast, in GL-CNT the largest ΔF/F₀ amplitudes occurred predominantly among regular contributors, although high-amplitude transients were also detected in highly active and sparsely active neurons (**Figure 3H,L**). These differences suggest that, beyond directly modulating neuronal activity, the surrounding microenvironment also shapes how neurons respond to the presence of CNTs. The association of stronger calcium signals with a regularly active neuronal population is particularly relevant to dopaminergic systems, in which sustained autonomous activity and recurrent calcium entry are closely linked to mitochondrial demand, oxidative stress, and neuronal vulnerability in Parkinson’s disease.[63] The GL-CNT phenotype may therefore provide a complementary microenvironment for investigating how sustained calcium activity interacts with these processes. Although calcium imaging does not directly measure membrane action potentials or synaptic connectivity, the broader distribution of calcium activity is consistent with enhanced functional recruitment within the engineered neuronal populations. Collectively, these observations indicate that calcium-transient amplitude is not strictly determined by event-frequency-based phenotype but instead represents a partially independent dimension of neuronal activity that can itself be modulated by the hydrogel microenvironment.

The activity phenotypes identified by clustering and the analysis of the raster plots, heatmaps, and mean activity profiles were further contextualized by the distributions of the coefficient of variation of the inter-event intervals (**CV**) (**Figure S5**). These broad distributions indicated heterogeneous temporal firing patterns that differed slightly among the SL-CNT, GL, and GL-CNT microenvironments. While most neurons exhibited moderately regular firing (CV ≈ 0.4– 0.7), distinct subpopulations of highly regular (CV < 0.3) and highly irregular (CV > 1) neurons were also observed in several hydrogel conditions. These findings were consistent with the functional heterogeneity observed in the raster plots. The relationship between event count and inter-event-interval CV showed no clear association, suggesting that increased neuronal activity was not necessarily accompanied by more regular firing. Thus, activity level and temporal regularity represented distinct features of the identified neuronal phenotypes (**Figure S5B,E,H**). Interestingly, consistent moderate inverse correlations between calcium-transient duration and CV were observed in SL-CNT, GL, and GL-CNT (**r = −0.35 to −0.41; Figure S5C,F,I**). Longer calcium transients therefore tended to occur in neurons with more regular inter-event timing. The recurrence of this relationship across the three analyzed conditions suggests that it represents a shared feature of these neuronal cultures rather than a relationship unique to one hydrogel microenvironment.

These analyses reveal that defined modifications of the hydrogel microenvironment did not merely alter the overall magnitude of neuronal activity but reshaped the distribution of functional states within the neuronal population. SL maintained a predominantly silent or sparsely active population, whereas the more organized GL microenvironment was associated with greater neuronal maturation, broader functional recruitment, and a dominant regular-contributor phenotype. CNT incorporation further expanded the range of neuronal behaviors, generating heterogeneous populations comprising highly active neurons, regular contributors, and a smaller proportion of sparsely active neurons with high-amplitude calcium transients manifesting in distinct neuronal clusters. These findings indicate that hydrogel composition and supramolecular organization can bias the emergence of distinct neuronal activity phenotypes within otherwise similarly differentiated neuronal populations, effectively turning microenvironmental design into a tuneability tool to study neurodevelopmental, neurometabolic and neurodegenerative diseases.

## Conclusions

By tailoring the synthetic conditions of chitosan hydrogels through controlled supramolecular organization and hydration states, tunable hydrogel microenvironments were successfully engineered. These hydrogels demonstrated excellent long-term stability under physiological conditions, making them promising candidates for *in vitro* neuronal culture platforms, including disease modeling applications. Their stability was primarily attributed to dynamic supramolecular crosslinking mediated by hydrogen bonding, with additional evidence suggesting possible contributions from π–hydrogen interactions and *in situ* crystallization of PABA. The resulting differences in supramolecular organization and water content generated distinct physicochemical microenvironments that markedly impacted NeuN and TUBB3 expression in human iPSC-derived dopaminergic neurons.

Immunostaining demonstrated that hydrogel supramolecular organization and CNT incorporation differentially influenced neuronal maturation and neurite network development, with the hydrated, highly organized GL microenvironment favoring higher NeuN levels and CNT-containing systems promoting TUBB3+ network formation. Importantly, computational analysis of spontaneous calcium activity revealed that these structural differences were accompanied by fundamentally distinct functional organizations of the neuronal populations. SL maintained a predominantly silent, less mature state, whereas GL favored a more uniformly recruited population dominated by regular contributors. CNT incorporation further diversified these functional states into strikingly similar functional profiles characterized by a predominance of highly active or bursting neurons, a smaller proportion of regular contributors, and only a minor population of sparsely active neurons. Additionally, CNT incorporation altered the relationship between calcium-event frequency and amplitude: SL-CNT supported high-amplitude transients among lower-event-count neurons, revealing a characteristic partial decoupling of amplitude and frequency, whereas GL-CNT concentrated the largest calcium responses predominantly within regular contributors. The microenvironments did not simply alter neuronal maturation or overall activity but reshaped how activity is distributed across neuronal phenotypes and how different dimensions of calcium signaling are coupled.

In conclusion, these findings demonstrate that engineered microenvironments bias neuronal phenotype, thereby providing a foundation for hierarchical control over complex *in vitro* brain and disease models.

## Supporting information

BIORXIV/2026/744416_Supplementary Material

## Author Contributions

BD designed, synthesized, and fully characterized the hydrogels. BD also performed confocal imaging analysis and fluorescence quantification of differentiated iPSCs, implemented and analyzed the computational calcium imaging workflow, and wrote the manuscript. LR performed the iPSC differentiation protocol and calcium imaging acquisition. PS contributed to hydrogel’s formulations testing and preliminary cytotoxicity analysis. GL performed the NMR experiment and its analysis. MP, NA, and SAM supervised the entire study, contributed to data interpretation, and critically revised/corrected the manuscript.

