## Supplementary material for "Tunable neuronal microenvironments drive distinct functional phenotypes in human iPSCs-derived dopaminergic neurons": BIORXIV/2026/744416_Supplementary Material

### Supporting Information

#### $^1$H NMR calibration and quantification of MA-functionalized chitosan

Maleic anhydride (MA) incorporation into chitosan was evaluated by $^1$H NMR using pyridine as an internal standard to determine whether varying the amount of MA reagent resulted in different degrees of maleoylation of MCS. The pyridine (Py) concentration was kept constant in all NMR samples, and its resonance at approximately 8.4 ppm was normalized to an integral value of 1. The vinyl proton signal of MA in the calibration solutions was integrated at approximately 6.15 ppm. In the modified chitosan samples, the corresponding resonance appeared between 5.95 and 6.05 ppm and was integrated accordingly. This chemical shift is consistent with changes in the local electronic environment following covalent incorporation of MA into the chitosan backbone.

The calibration curve was constructed using four MA concentrations in the presence of a constant pyridine concentration (2.6 mg·mL$^{-1}$). MA concentration was expressed in mmol·mL$^{-1}$ and calculated using the molecular weight of MA (98.06 g·mol$^{-1}$; 98.06 mg·mmol$^{-1}$). Linear regression of the MA/Py integral ratio against MA concentration was described by:

$$\frac{I_{\mathrm{MA}}}{I_{\mathrm{Py}}} = 3.7203\, C_{\mathrm{MA}} - 0.16075 \qquad \text{Eq}(\mathbf{8})$$

and therefore:

$$C_{\mathrm{MA}} = \frac{(I_{\mathrm{MA}}/I_{\mathrm{Py}})+0.16075}{3.7203} \qquad \text{Eq}(\mathbf{9})$$

where $C_{MA}$ is the estimated concentration of incorporated MA (mmol·mL$^{-1}$), and $I_{MA}$ and $I_{Py}$ are the integrated areas of the MA vinyl and pyridine resonances, respectively.

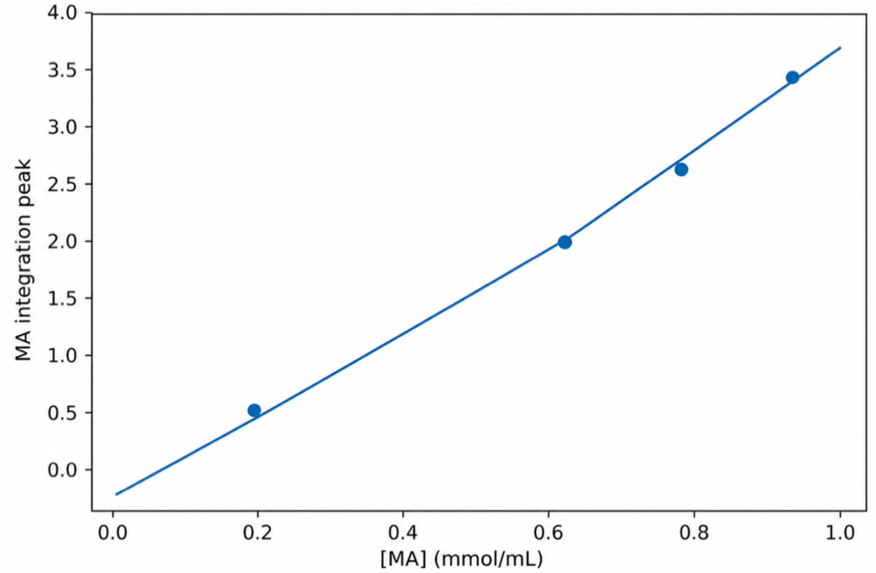


**Figure S1. $^1$H NMR calibration curve for the quantification of MA-substitution on chitosan.** A total of four concentrations of maleic anhydride (MA) was used in the presence of constant amount of pyridine.

Functionalized MCS samples were prepared at approximately 10 mg·mL$^{-1}$ MCS concentration (6 mg in 600 µL). In both MCS samples prepared using 3.5 g and 7 g of MA, the MA/maleoyl-related integral in the 5.95–6.05 ppm region was approximately 0.01 after normalization of the pyridine signal to 1. This corresponded to an estimated maleoyl content of approximately 4.6 mmol·g$^{-1}$. Because the sample signals fell below the lowest concentration included in the calibration curve, these values were obtained by extrapolation and should therefore be considered estimates rather than precise absolute quantifications. Nevertheless, the comparable signals obtained for the two samples indicate that increasing the MA amount from 3.5 to 7 g produced no detectable increase in maleoylation under these conditions, supporting the use of 3.5 g MA for subsequent experiments.

**Table S1.** Quantification of maleoylation content upon changing the concentration of maleic anhydride.

| Sample | $I_{Py}$ | $I_{MA}$ | $C_{MA}$(mmol mL$^{-1}$) | MA content (mmol g$^{-1}$) |
|---|---|---|---|---|
| MCS–3.5 g MA | 1.000 | 0.01 | 0.04590 | 4.590 |
| MCS–7 g MA | 1.000 | 0.01 | 0.04590 | 4.590 |

Py, pyridine internal standard; $I_{Py}$, integrated signal of pyridine normalized to 1; $I_{MA}$, integrated signal assigned to the maleoyl group; $C_{MA}$, MA concentration determined from the calibration curve using the corresponding NMR integration.

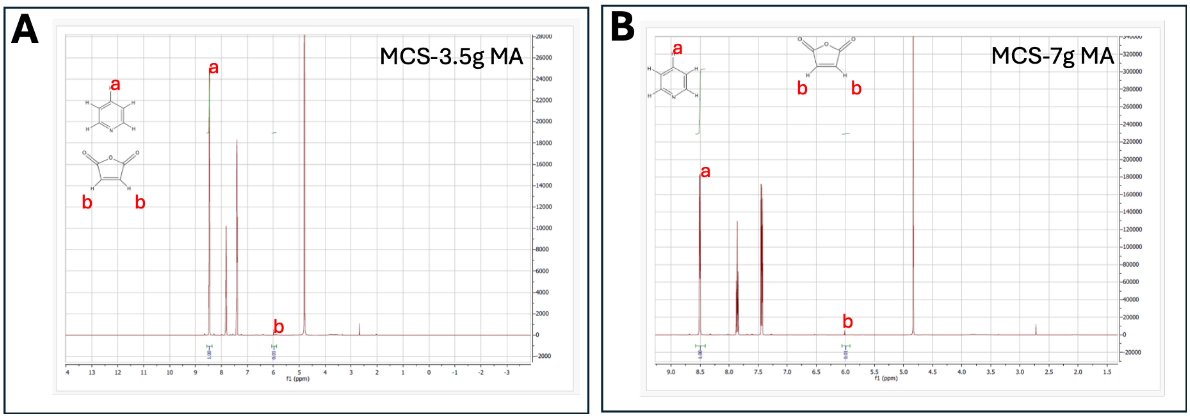


**Figure S2. $^1$H NMR of MCS samples with different maleic anhydride concentrations.** (**A**) $^1$H NMR of MCS prepared from 3.5g of maleic anhydride (MA) *vs.* (**B**) MCS prepared from 7g of MA.

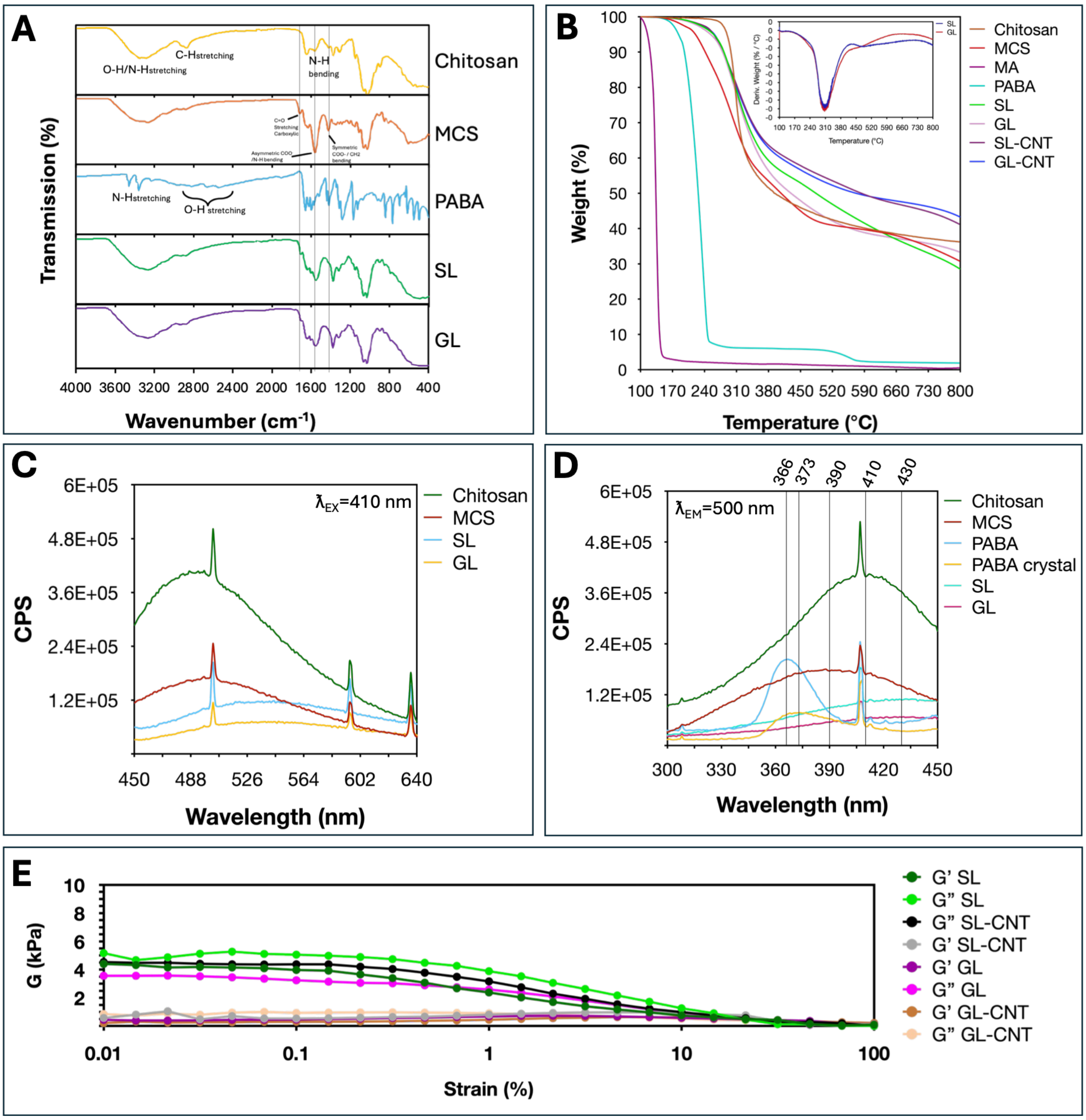


**Figure S3. Physicochemical and rheological characterization of the hydrogel systems.** (**A**) Fourier transform infrared (FTIR) spectra of chitosan, 4-Amino benzoic acid (PABA), maleated chitosan (MCS), SL, and GL, confirming the chemical composition of the hydrogel systems. (**B**) Thermogravimetric analysis (TGA) profiles of the corresponding samples. (**C**) Fluorescence emission spectra in counts per second (CPS) of chitosan, MCS, SL, and GL measured at an excitation wavelength of 410 nm. (**D**) Fluorescence excitation spectra of chitosan, MCS, PABA (microcrystalline), PABA crystals obtained by solvothermal method, SL, and GL recorded at an emission wavelength of 500 nm. (**E**) Oscillatory amplitude sweeps measurements showing the linear viscoelastic region (LVR) of the hydrogel formulations. G', storage modulus. G'', loss modulus. CNT, carbon nanotubes.

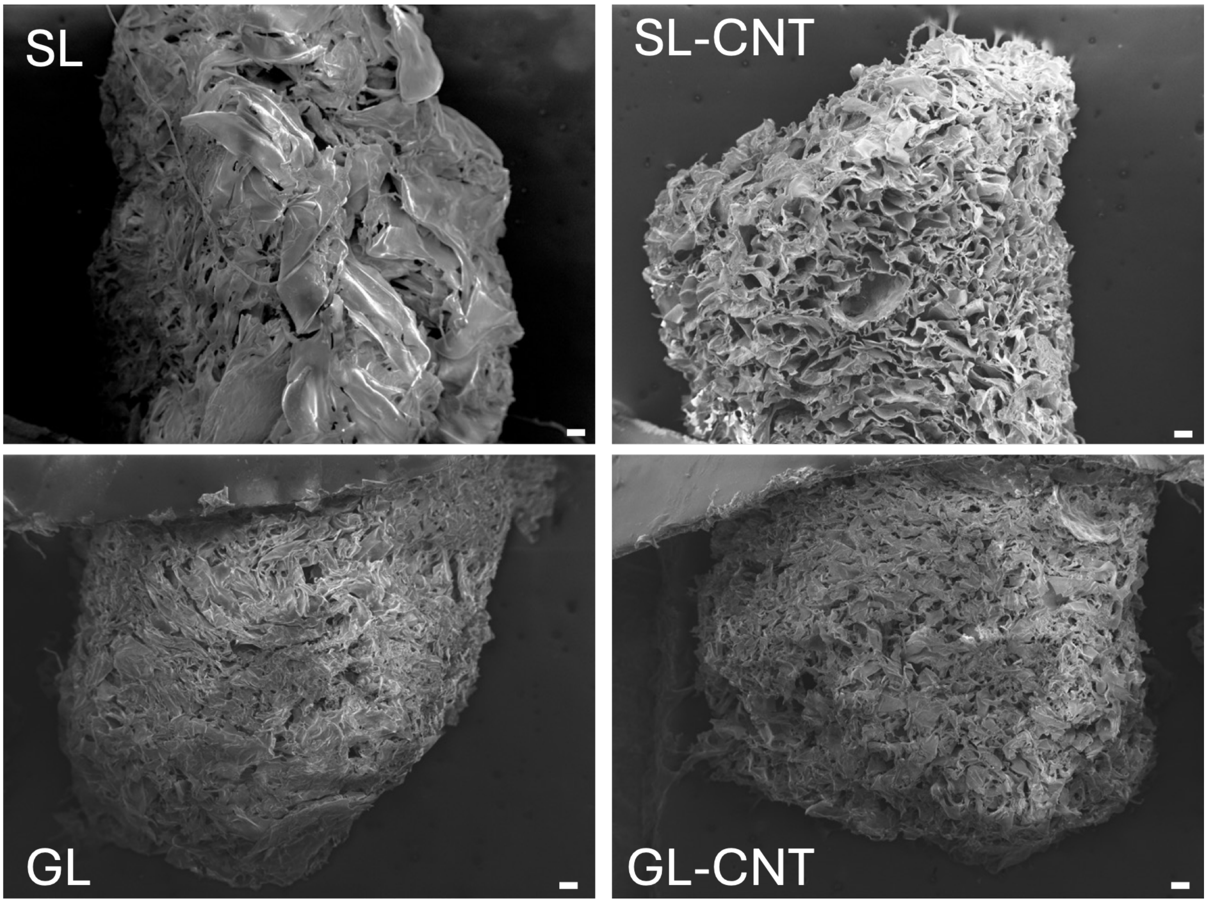


**Figure S4. Scanning Electron Microscopy (SEM) images of chitosan hydrogels.** SEM micrographs showing the overall morphology and porous microstructure of the hydrogels. The corresponding hydrogel formulation is indicated on each image. Scale bars, 100 µm.

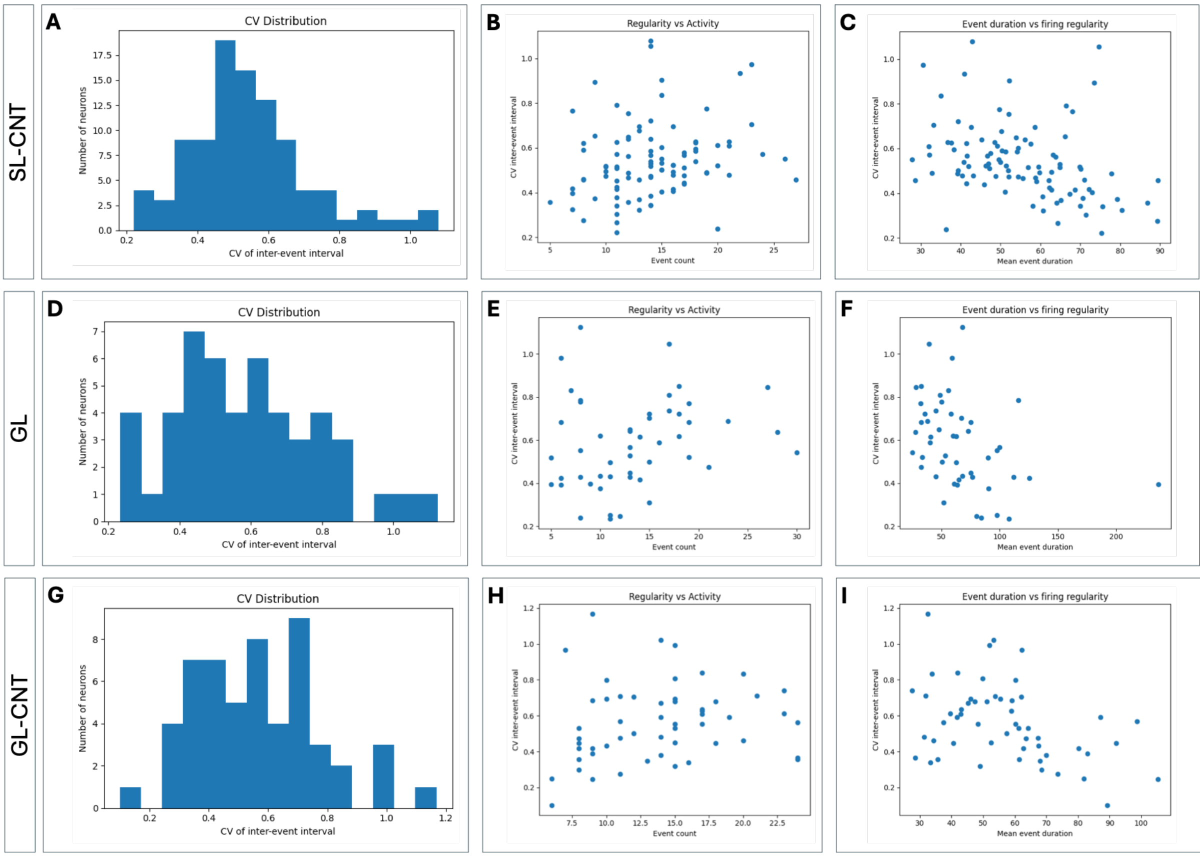


**Figure S5. Firing and activity variability analysis across hydrogel formulations.** (**A-C**) Functional activity analysis of neurons cultured in the SL-CNT hydrogel. (**A**) Distribution of the coefficient of variation (CV) of inter-event intervals, describing the variability of neuronal firing. (**B**) Relationship between firing regularity (CV) and neuronal activity (event count). (**C**) Relationship between calcium transient duration and firing regularity (CV). (**D-F**) Corresponding analyses for the GL hydrogel. (**G-I**) Corresponding analyses for the GL-CNT hydrogel. Each data point represents an individual neuron.
